# A subcortical origin for rapid, target-oriented corticospinal excitability modulation during visually guided reaching

**DOI:** 10.1101/2022.12.13.520229

**Authors:** Rechu Divakar, Gerald E. Loeb, Brian D. Corneil, Guy Wallis, Timothy J. Carroll

## Abstract

During visually guided reaching, proximal limb muscles can be activated within 80 ms of target appearance. Such “express” visuomotor responses are temporally aligned with target appearance rather than movement onset, and invariably tuned towards the direction of the visual target regardless of the instructed reach direction. These features prompt the hypothesis that express visuomotor responses are driven by a subcortical pathway. We tested this by measuring the changes in Motor Evoked Potential (MEP) size following Transcranial Magnetic Stimulation (TMS) or Transcranial Electrical Stimulation (TES) of the motor cortex, as participants reached either towards or away from visual targets. We found that 70-80 ms after target presentation, MEPs in a primary shoulder flexor muscle (pectoralis major) were oriented towards the target direction regardless of whether the participant subsequently reached towards or away from the target. Similar target-oriented MEP modulations occurred in posterior deltoid and biceps brachii muscles, whereas MEPs in a finger muscle were affected neither by target nor reach direction. Critically, there were no significant differences in modulation of responses to TMS versus TES across all reaching conditions, which suggests that the target-oriented modulation occurs downstream of the motor cortex output neurons. Combined, our results are consistent with a subcortical rather than cortical origin for the earliest changes in corticospinal excitability following visual target onset. A prime candidate for such subcortical modulation includes the superior colliculus and reticular formation.

## INTRODUCTION

The control of visually guided reaching in humans has long been attributed to a sensory to motor transformation that relies on processing within the visual, parietal and motor cortices (Kalaska et al., 1997; Kalaska, 2009; Filimon, 2010; Vesia and Crawford, 2012; Omrani et al., 2017). However, recent observations in experiments that require immediate limb responses have prompted consideration of alternative neural control networks for reaching. When humans reach to a visual target, proximal muscles can be activated within 80 ms of the target presentation. These express visuomotor responses share phenomenological similarities with express saccades, which are rapid eye movements executed in the direction of a visual target triggered by the first arrival of stimulus-related visual information in the superior colliculus. Express saccades are almost exclusively directed toward the location of the visual target, even if they occur during an “anti-saccade” task in which the subject is instructed to look elsewhere (Everling et al., 1999). Express responses in shoulder muscles are also tuned to bring the limb toward a visual target regardless of task instructions or the ultimate direction of limb motion (Gu et al., 2016; Contemori et al., 2022a). The onset latencies of express saccades and express visuomotor responses are similar; distributed around 100 ms after the target presentation (Fischer and Ramsperger, 1984; Kingstone and Klein, 1993; Kozak et al., 2019; Contemori et al., 2021). These phenomenological similarities between express saccades and express visuomotor responses have led many authors to speculate that these phenomena share neural substrates (Boehnke and Munoz, 2008; Corneil and Munoz, 2014). Given that the neural pathway responsible for generating express saccades is known to involve the midbrain superior colliculus and the brainstem reticular formation (Munoz et al., 2000), a tecto-reticulo-spinal origin for express visuomotor responses in the muscles that produce reaching seems plausible.

The short latency of the express visuomotor response is consistent with the transmission time from the retina to the muscle through a tectoreticulospinal pathway, and represents a key argument in favour of a subcortical origin for express responses. However, on latency grounds alone, it remains possible that express responses could be generated within the cerebral cortex. Indeed, fast cortical pathways that carry visual information to motor areas are well documented in primates. In simple visuomotor tasks like hand-lifting a lever in response to a visual stimulus (Gemba et al., 1981), or hand-releasing a lever to perform a visual discrimination task (Ledberg et al., 2007), local field potentials have been recorded from the motor cortex of monkeys at 40-60 ms after visual stimulus onset. In a task where monkeys had to align a cursor over a vertical target line by wrist flexion or extension, target-related spike activity was observed in the motor cortex around 40 ms after the appearance of the visual stimuli (Wong et al., 1982; Kwan et al., 1985). Similar responses were reported at around 60 ms after presentation of visual stimuli to prompt elbow flexion or extension movements (Lamarre et al., 1981, 1983). In choice visuomotor tasks involving reaching to multiple visual targets, unit recordings showed directionally specific responses in the premotor and primary motor cortices of monkeys as early as 50 ms (Murphy et al., 1982; Georgopoulos et al., 1989; Cisek and Kalaska, 2005; Reimer and Hatsopoulos, 2010; Ames et al., 2014). Anti-reach paradigms that dissociate the visual stimulus and response directions showed that by around 70 ms, the early motor cortical responses were tuned to the visual stimulus location rather than the movement direction (Crammond and Kalaska, 1994; Riehle et al., 1997). These observations are consistent with the idea that early motor cortex responses are related to stimulus-encoding, as opposed to goal-related movement preparation and execution, so such activity could be a source of motor output that leads to express visuomotor responses.

The rapid and stimulus-directed features of early motor cortex activity in non-human primates are similar to those of express visuomotor responses in humans. However, while these studies support the existence of fast cortical transmission of visual signals to motor cortical areas in non-human primates, it is important to acknowledge that the response latencies observed in non-human primates are not directly applicable to humans. While there is currently no direct evidence to show the timing of stimulus related visual inputs to the motor cortex in humans, it is known that the timings of both express saccades and express muscle responses can be ∼20 ms faster in monkeys compared to humans (Fischer, 1986; Goonetilleke et al., 2015; Cecala et al., 2023).

Non-invasive brain stimulation provides a means to probe the responsiveness of cortical and subcortical elements of the corticospinal tract in humans during tasks that can elicit express visuomotor responses. If the muscle responses to brain stimulation are modulated systematically by the location of a visual target, it would indicate that the circuits responsible for the evoked responses have access to target-related information at the time of stimulation. Thus, it should be possible to chart the time-course of target-related visuomotor processing within the human corticospinal pathways using non-invasive brain stimulation. Descending signals responsible for express visuomotor responses must eventually arrive at and drive action potentials in spinal neurons. Given this, any experimental probe of corticospinal tract excitability will be subject to the excitability state of spinal interneurons and motoneurons. The site of excitability modulation due to express visuomotor response processing can be differentiated through comparison of responses to Transcranial Electrical Stimulation (TES) and Transcranial Magnetic Stimulation (TMS) because they generate electric fields that are oriented orthogonally within the cerebral cortex (Ruohonen and Ilmoniemi, 2005). At threshold intensities, TMS activates the corticospinal neurons indirectly via horizontally oriented mono- and oligosynaptic inputs, whereas TES activates the vertically oriented corticospinal pyramidal neurons directly (Rothwell, 1991; Edgley et al., 1997; Di Lazzaro et al., 1998a; Rothwell, 2005; Siebner et al., 2022). Therefore, at threshold intensities, muscle responses evoked by TMS depend on the excitability of cortical interneurons with mono- and oligosynaptic connections to the corticospinal output neurons and the corticospinal neurons themselves. By contrast, the responses to threshold-level TES should not be influenced by the excitability of neurons in the motor cortex, but rather only by downstream elements of the corticospinal pathways. The fact that voluntary contraction, which affects cortical excitability, influences the magnitude of descending spinal volleys generated by TMS but not TES also suggests that TES evoked responses originate from direct excitation of corticospinal axons (Di Lazzaro et al., 1998b, 1998a, 1999).

If express visuomotor responses are generated cortically, there should be significant differences in the time course and/or magnitude of the target-related modulation of TMS and TES responses. Firstly, target-related excitability changes would be expected to arise earlier in TMS compared to TES, as there should be intracortical responsiveness changes prior to release of descending corticospinal outputs. Secondly, the overall change in MEPs evoked by TMS should be greater, as the MEPs should be influenced by both subcortical and cortical responsiveness changes, whereas TES responses should only be modulated subcortically. Conversely, if express visuomotor responses are generated sub-cortically, the time-course and magnitude of evoked responses to both forms of brain stimulation should be similar, as both would be affected only by spinal level modulation. Such comparisons between TMS and TES responses have been made previously to argue that sensory-driven modulations in corticospinal excitability during visually guided reaching were at least partially mediated by subcortical mechanisms (Suzuki et al. 2021).

Using this logic, we sought to test the origins of express visuomotor responses by asking human participants to make reaching movements in response to visual targets while we applied non-invasive brain stimulation at various times after target presentation. On a given trial, participants performed either a pro-reach, where they reached towards the direction of the visual target, or an anti-reach, where they reached away from the direction of the visual target. In two experiments, we systematically varied the latency of corticospinal tract stimulation between target presentation and movement initiation and tracked the changes in motor evoked potential amplitude in proximal limb muscles as a function of time. In the first experiment, using only TMS, we tested whether the early corticospinal excitability changes following visual target presentation were biased in the direction of the visual target or the reach goal. In the second experiment we compared TES and TMS responses at various latencies to identify whether changes in corticospinal excitability occurred at cortical or subcortical sites within the corticospinal tract.

## RESULTS

### Experiment 1 – Corticospinal excitability is biased by the target location 70 – 80 ms after visual target presentation in functionally relevant muscles

In experiment one, we sought to establish when corticospinal excitability was first directionally biased after target presentation prior to visually guided reaching. To this end, we stimulated the motor cortex using transcranial magnetic stimulation (TMS) to elicit motor evoked potentials from the pectoralis major, posterior deltoid, biceps brachii and first dorsal interosseous muscles. On a given trial, participants either reached towards (pro-reach) or away (anti-reach) from a visual target, as a single magnetic pulse was randomly given at 60, 70, 80, 90 or 100 ms after the onset of the visual target. Stimulation was also given just prior to the onset of the reach; this stimulation latency was called the *voluntary* latency. To see whether the early corticospinal excitability following target onset was biased in the direction of the visual target or the reach goal we tracked changes in MEP size as a function of stimulation latency for two reach types (pro- and anti-reach) and two target directions (left and right).

First, we looked at how participants performed in the task. Participants generally made rapid reaches to the nominated direction in both pro and anti-reach conditions (reaction time: pro-reach = 227 ± 28 ms, anti-reach = 254 ± 32 ms). As expected, the requirement to reach away from the physical location of the target delayed the initiation of limb movement (two-way RM ANOVA; main effect of reach-type: *F*_1,19_ = 34.5, *P* < 0.001; **Figure 1A**). However, the location of the target did not significantly affect the reaction time (main effect of target direction: *F*_1,19_ = 0.5, *P* = 0.482; interaction: *F*_1,19_ = 1.9, *P* = 0.184; **Figure 1A**). The occurrence of reach errors, where participants did not reach according to task instruction, was significantly more common for anti-than pro-reaches (*t*_19_ = 3.2, *P* = 0.004; **Figure 1B**). The target location did not significantly affect the number of reach errors (*t*_19_ = 0.5, *P* = 0.586; **Figure 1B**). Next, we combined the data from both pro- and anti-reach trials and examined the effect of brain stimulation on task performance. We found that participants made more errors during trials with TMS than in trials without TMS (*t*_19_ = 5.3, *P* < 0.001; **Figure 1D**), while their reaction times did not differ significantly between the two conditions (*t*_19_ = 1.7, *P* = 0.106; **Figure 1C**). Next, we assessed whether participants had overt express visuomotor responses according to standard criteria applied in previous studies. We ran a receiver operator characteristic (ROC) analysis on the *baseline* reaching blocks where no brain stimulation was given (see Methods: Express Visuomotor Response Detection). Only 2 out of the 20 participants (p1 and p4) had an express visuomotor response according to standard criteria, and only in the pro-reach condition and not in the anti-reach condition. This was surprising as previous studies using similar paradigms have reported more prevalent express visuomotor responses. The reasons for this discrepancy are unclear, but may include anticipation of the brain stimulation or the use of large circular EMG electrodes, rather than narrow bar electrodes with 10mm inter-electrode distance.

**Figure 1.**
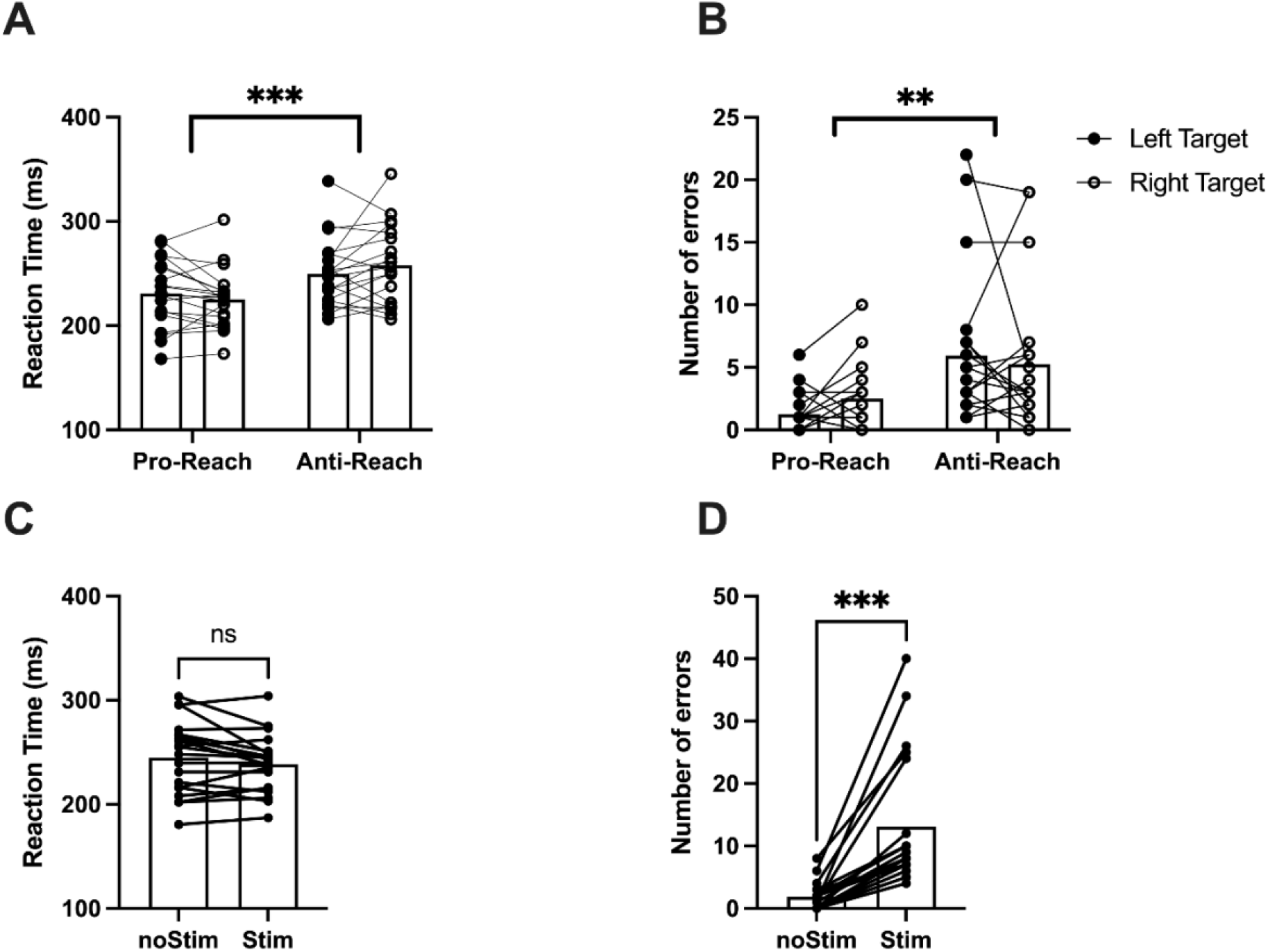
***A***: Reaction time of correct reaches in the pro-reach and anti-reach condition. ***B***: Number of reach errors in pro- and anti-reach condition. ***C***: Reaction time in trials with and without brain stimulation. ***D***: Number of reach errors in trials with and without brain stimulation. Error bars show 95% CI. *P < 0.05, **P < 0.01, ***P < 0.001.

**Figure 2** shows an exemplar participant (p3) who was classified negative for express visuomotor responses. Nonetheless, single-trial EMG analysis (see Methods) and the average EMG trace show that this participant had some trials with EMG activity onsets in the 70-120 ms range, typically considered as the express response window (**Figure 2A-F**). Box plots show that the distribution of EMG onset times include trials with onsets within the express response time-window for all participants (**Figure 2G**). The presence of voluntary muscle activity in the time window for application of brain stimulation would amplify the descending volleys evoked by brain stimulation due to increased motoneuronal activation. Thus, the time course of voluntary EMG in control trials during reaching blocks with brain stimulation is documented in **Figure 5**, and incorporated into MEP interpretation.

**Figure 2.**
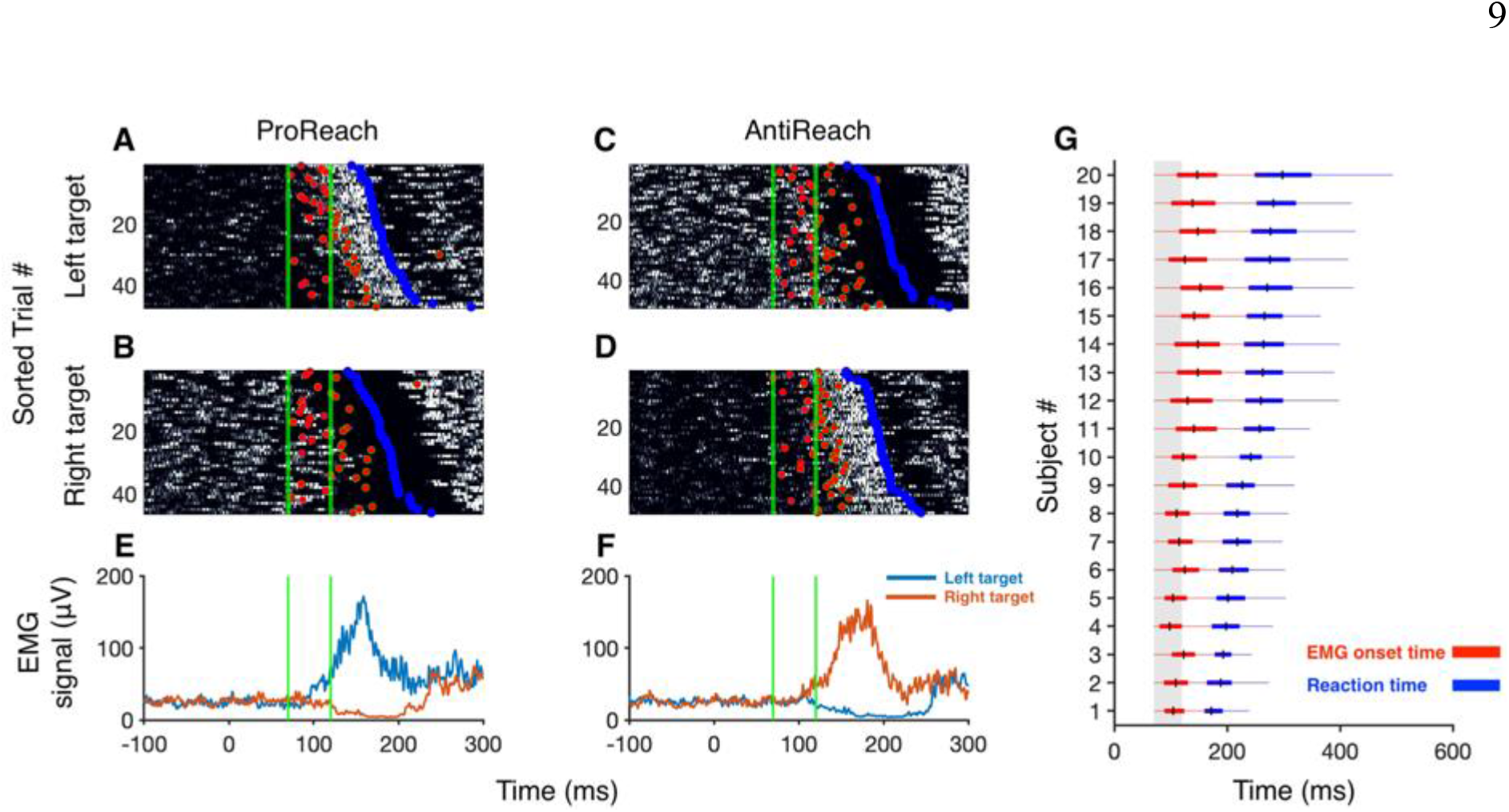
***A-D***: Exemplar participant’s (participant number 3 in ***G***) raster plots showing the grey-scale coded EMG activity in individual trials across the pro-reach (***A&B***) and anti-reach (***C&D***) conditions for two target locations. White regions in the raster plot represent an increase in EMG and dark regions represent a decrease in EMG. Trials in every condition were sorted based on the reaction time (blue dots), the trial number is shown on the y-axis. The x-axis shows the time (in ms) with respect to the target onset. Red dots represent the EMG onset or offset detected using the single-trial EMG activity detection algorithm (see: Methods). ***E&F:*** Average EMG (in μV) in pro-reach and anti-reach condition respectively. The green lines represent the express visuomotor response time window (70-120 ms). ***G:*** Boxplots showing the distribution of the onset of EMG activity (red boxplot) and reaction time (blue boxplot) for all 20 subjects. The box represents the interquartile range, the black line in the box plot represents the median reaction time, and the whiskers represent the minimum and maximum. Boxplots are sorted based on the median reaction time of the participants. The gray shaded regions show the express visuomotor response time window (70-120 ms).

In the first experiment, our main variable of interest was the amplitude of motor evoked potentials (MEPs) in the pectoralis major muscle resulting from transcranial magnetic stimulation (TMS). **Figure 3A&B** depicts the raw MEPs obtained from a representative participant, with the mean traces superimposed on the individual waveforms. **Figure 3C&D** shows the average MEP amplitude at each stimulus latency for this participant, and illustrates that MEP amplitudes are greater for leftward than rightward targets from ∼ 70ms after target presentation in the pro-reach condition. Given that the Pectoralis acts to move the hand leftward in this posture, the data show that responses to brain stimulation in this person were biased towards the visual target direction, which also coincides with the direction of the reach in this condition. Crucially, in the anti-reach condition, where the visual target direction is opposite to the reach direction, there was also a small bias towards greater MEP amplitudes for the left target at around 80-90 ms. This suggests that early responses to brain stimulation were biased toward the visual target direction rather than the direction of hand motion in this participant.

**Figure 3.**
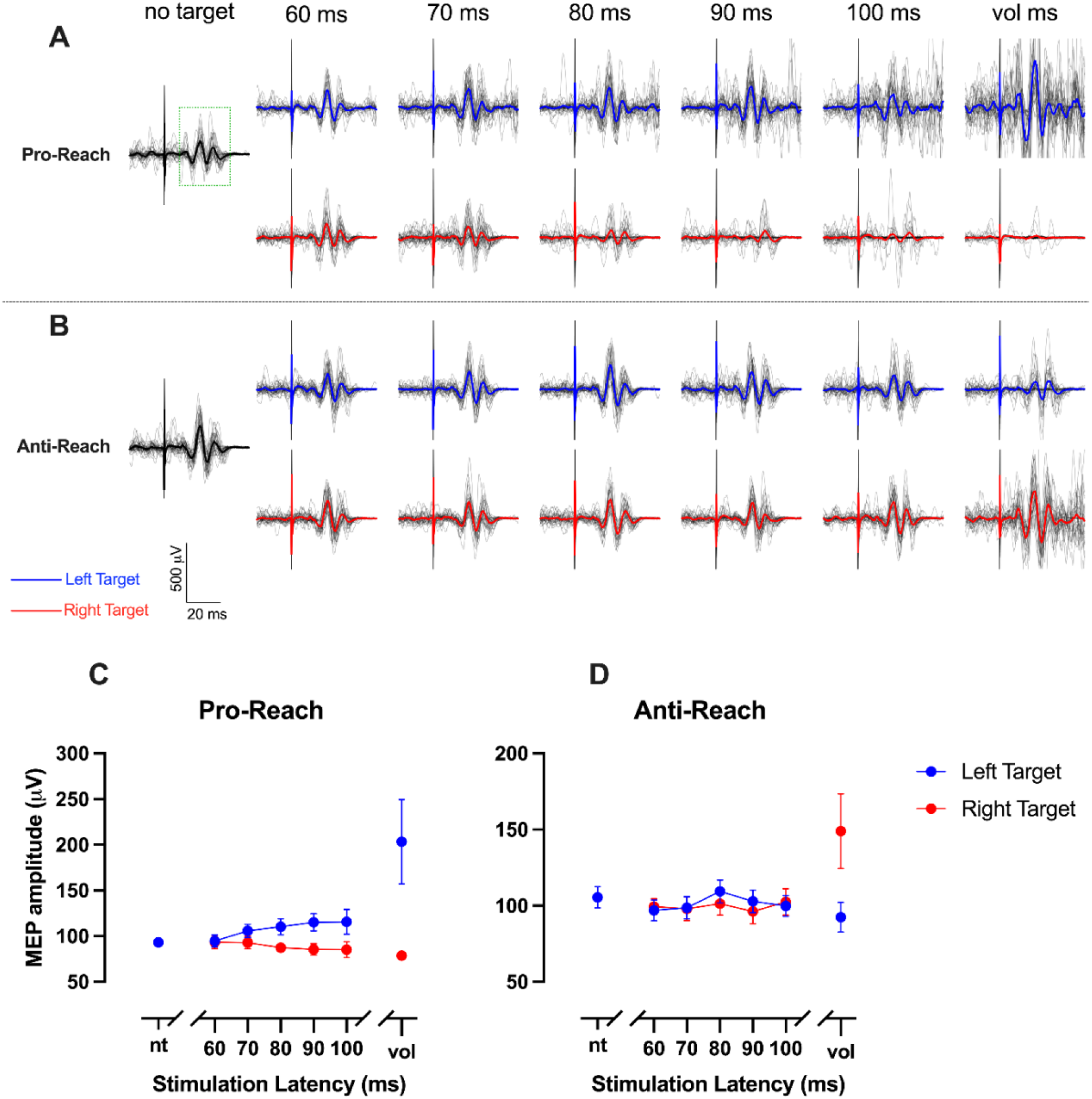
Motor evoked potentials recorded from the pectoralis major muscle of an exemplar participant. Individual MEP traces from each stimulus timing with the mean MEP overlaid are shown for pro-reach (***A***) and anti-reach (***B***) conditions. The green dotted box in the *no target* condition for Pro-Reaches shows the time-window that was visually identified for calculating the MEP amplitude. Time course of the average MEP amplitude for the same participant in the pro-reach (***A***) and anti-reach (***B***) conditions. nt – no target, vol – voluntary latency. Error bars show 95% confidence intervals.

**Figure 4A&B** shows that the results from this exemplar participant were broadly similar across our sample of 20 people. To verify this, we used a group-level repeated-measures ANOVA with target direction (Left and Right) and stimulation latency (60, 70, 80, 90, 100, and *voluntary*) as factors. Separate ANOVAs were run for pro- and anti-reach conditions. MEP size varied significantly with visual target direction and the time of TMS delivery according to a significant interaction between target direction and stimulation latency for both pro- (*F*_1.5,28.4_ = 15.3, *P* < 0.001) and anti-reaches (*F*_1.4,26.2_ = 25.6, *P* < 0.001). MEPs elicited by stimulation at the *voluntary* latency were much larger and more variable across participants than MEPs evoked by stimulation at the fixed latencies. To ensure that the interaction effects were not driven predominantly by the large MEPs at the *voluntary* latency, we reran the ANOVAs after excluding the *voluntary* latency (see **Figure 5A&B)**. When considering only the fixed latencies between 60-100 ms, a significant target direction by stimulation latency interaction remained in both pro-reach (*F*_1.5,29.3_ = 6.2, *P* = 0.010) and anti-reach (*F*_2.2,41.2_ = 16.5, *P* < 0.001) conditions.

**Figure 4.**
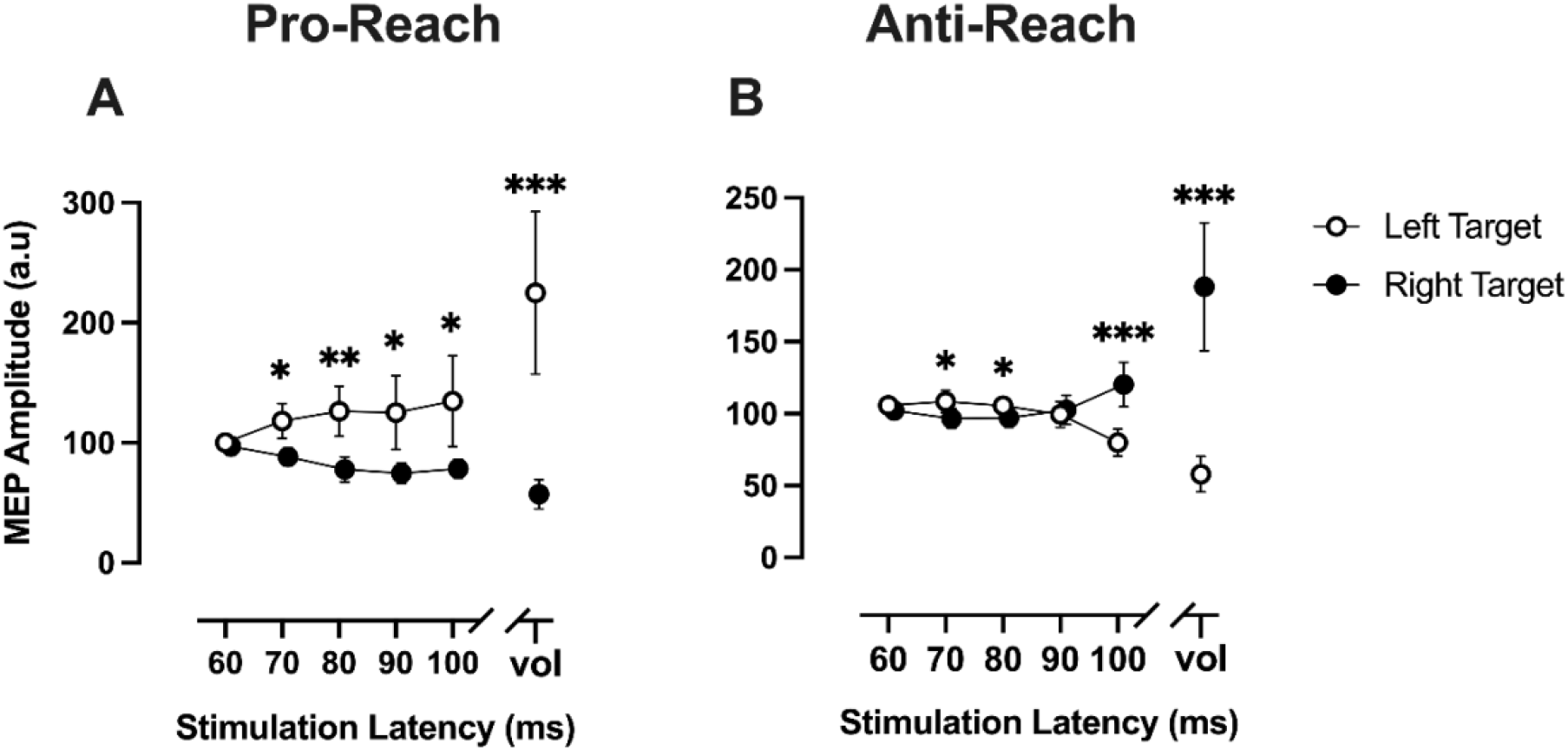
***A & B***: Time course of MEP amplitude as a function of stimulation latency. First column shows results in the pro-reach condition and second column shows results in the anti-reach condition. Black circles represent the left target direction and white circles represent the right target direction. Error bars show 95% CI. *P < 0.05, **P < 0.01, ***P < 0.001.

**Figure 5.**
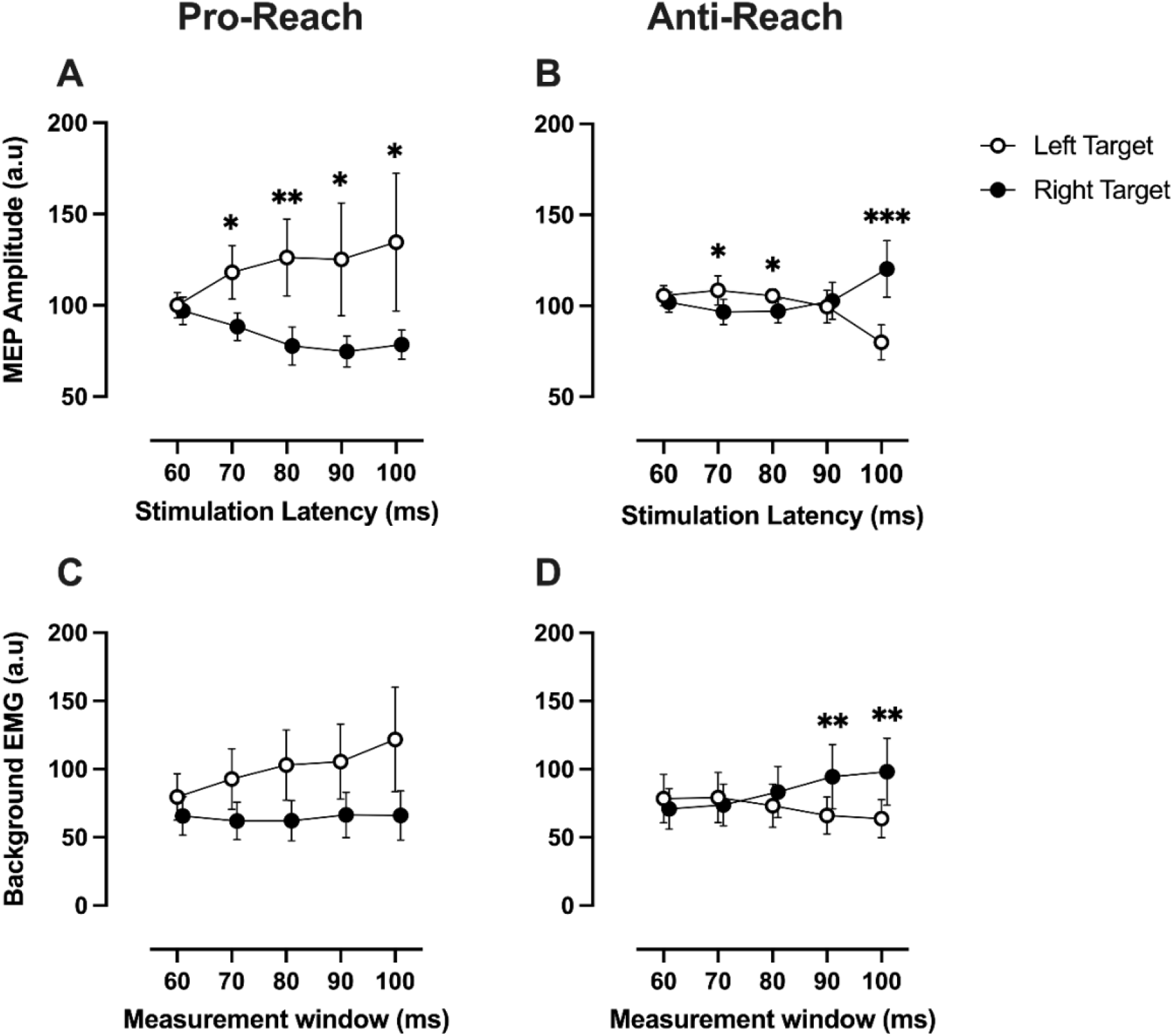
***A & B***: Time course of MEP amplitude as a function of stimulation latency. ***C & D***: Time course of background EMG as a function of measurement window. First column shows results in the pro-reach condition and second column shows results in the anti-reach condition. Black circles represent the left target direction and white circles represent the right target direction. Error bars show 95% CI. *P < 0.05, **P < 0.01, ***P < 0.001.

To identify when the location of target significantly influenced MEP size, we used Holm-Šídák multiple comparisons to test the difference between the left and right target MEPs at each stimulation latency. The target direction did not significantly influence the excitability of the corticospinal tract at 60 ms in either pro- or anti-reach conditions (**Figure 5A&B**), as the MEPs for the left target did not differ significantly from that of the right target (pro-reach: *P* = 0.584, anti-reach: *P* = 0.329). Crucially, MEP sizes at 70 and 80 ms were biased by the location of the visual target. Regardless of the direction in which the participants reached, the MEPs for the left target was significantly larger than the right target at 70 ms (pro-reach: *P* = 0.019, anti-reach: *P* = 0.036) and 80 ms (pro-reach: *P* = 0.008, anti-reach: *P* = 0.036). This result is expected in pro-reaches, as the target and movement goal are in the same direction, hence MEPs were enhanced or inhibited when PEC served as an agonist or antagonist, respectively (**Figure 5A**).

Critically, we observed a similar result for the anti-reach condition (**Figure 5B**), which shows that despite reaching in the direction opposite from the target, the corticospinal excitability at 70 and 80 ms was tuned to bring the hand towards the target location. In the pro-reach condition, the left target MEPs remained larger than the right target MEPs at 90 ms (*P* = 0.021) and 100 ms (*P* = 0.021) (**Figure 5A**). However, in the anti-reach condition the MEPs converged and overlapped at 90 ms (*P* = 0.612) and flipped in the opposite direction at 100 ms such that now right target MEPs were larger than the left target MEPs (*P* < 0.001). Thus, by 100 ms, the amplitudes of MEPs were higher when a leftward reach was the motor goal compared to when a rightward reach was the motor goal (**Figure 5B**). From this time onward, the evoked response in the PEC muscle was facilitated if its activity would move the hand towards the motor goal and inhibited if its activity would move the hand away from the motor goal. When the stimulation was delivered at the voluntary latency, which had distinct timings for pro-and anti-reaches in each participant, (average voluntary stimulation latency for pro-reach: 143 ± 28.83 ms, range = [110 220]; anti-reach: 167 ± 27.22 ms, range = [130 240]) MEPs induced in the PEC muscle were strongly tuned towards the motor goal. In both pro- and anti-reach conditions, MEPs were larger when a left reach was the motor goal (pro-reach: *P* < 0.001; anti-reach: *P* < 0.001; **Figure 4**). The data indicate that around 70-80 ms after target presentation the excitability of the corticospinal tract is biased in the direction of the visual target. But from 100 ms onwards the bias is consistent with the direction of the reach goal.

To investigate this further, we looked at the muscle activity in the no-stimulation trials (*noStim* trials) that were intermixed with the stimulation trials. Since these trials were randomly intermixed, they can inform us about the muscle activity during task performance in the absence of TMS. For each participant and each stimulation latency, we measured the average of the rectified EMG from the same ‘measurement time window’ that was used to calculate the MEP in the stimulation trials. This window was unique for each participant as the onset and shape of the MEP varied across participants. The EMG amplitude measured this way was expressed as a percentage of the muscle activity in the *noTarget* trials in a 25 ms window before stimulation onset. **Figure 5C&D** shows the normalized muscle activity in the MEP time windows in the absence of magnetic stimulation. To check how the background muscle activity changed in the early latencies at which we observed target-driven MEP changes, we used a RM-ANOVA with measurement window and target direction as the repeated measures. We excluded the *voluntary* measurement window from the analysis of variance as we were interested only in the early muscle activity. In the pro-reach condition, there was increased muscle activation across the measurement window for the left target, but the muscle activity remained relatively consistent for the right target (**Figure 5C**). The RM-ANOVA results supported these conclusions, there were significant main effects of measurement window (*F*_1.3,32.3_ = 10.1, *P* = 0.002) and target direction (*F*_1, 19_ = 12.7, *P* = 0.002). The interaction effect was not significant, but marginal (*F*_1.1,21.9_ = 3.8, *P* = 0.058). In the anti-reach condition, the background activity remained relatively similar for the left and right targets until 80 ms, but from 90 ms onward the muscle activity increased in manner to facilitate the limb movement towards the target (**Figure 5D**). This conclusion was supported by the RM-ANOVA which showed a significant interaction between the target direction and measurement window (*F*_1.3,25.4_ = 12.1, *P* < 0.001) and significant differences at 90 and 100 ms via post-hoc tests (90 and 100 ms, *P* < 0.01). Thus, in both pro- and anti-reach conditions, voluntary EMG was modulated in the time-windows in which we observed target-driven MEP changes. The early changes in background EMG correspond broadly with the timing of express visuomotor responses, despite the fact that only two participants were positive for express visuomotor responses as defined by standard criteria. It is clear that the MEP modulations we observed were influenced by the changes in early motoneuron activation reflected by the EMG data. However, if motoneuron activation were the sole cause of MEP modulation, then the time course of MEP responses should perfectly match the EMG responses. As the patterns of EMG and MEP modulations differ somewhat, it suggests that TMS induced MEPs (**Figure 5A&B**) did not merely reflect motoneuron activation, but were also influenced by the state of additional components of the corticospinal networks during the early stages of reaching. It appears that TMS is a powerful method to study the time course of neural events that underlie express visuomotor responses, especially when the express responses are weak.

Next, we checked whether other arm and hand muscles showed early target-driven corticospinal excitability changes like those seen in the PEC. **Figure 6** shows the motor evoked potentials from the biceps brachii (BIC), posterior deltoid (PD) and first dorsal interosseous (FDI). Biceps brachii muscle responses essentially replicated the results from the PEC, apart from a slight delay in time course of MEP size modulations that might relate to axonal conduction times. The location of the visual target and stimulation latency strongly influenced the MEPs at the early latencies, in both pro-reach (main effect of the target direction: *F*_1,19_ = 27.3, *P* < 0.001; main effect of stimulation latency: *F*_1.7,32.1_ = 7.0, *P* = 0.004; interaction: *F*_1.7,31.8_ = 15.6, *P* < 0.001) and anti-reach conditions (main effect of the target direction: *F*_1,19_ = 1.4, *P* = 0.255; main effect of stimulation latency: *F*_1.7,32.7_ = 3.5, *P* = 0.045; interaction: *F*_3.2,60.8_ = 8.2, *P* < 0.001). In the pro-reach condition, the MEP amplitude was larger for the left target compared to the right from 80 ms onwards (for 80, 90 and 100 ms: *P* < 0.001, **Figure 6A)**, whereas in the anti-reach condition, MEP amplitudes were representative of the target direction at 80 ms (*P* = 0.048) but flipped at 100 ms (*P* = 0.015) (**Figure 6B**).

**Figure 6.**
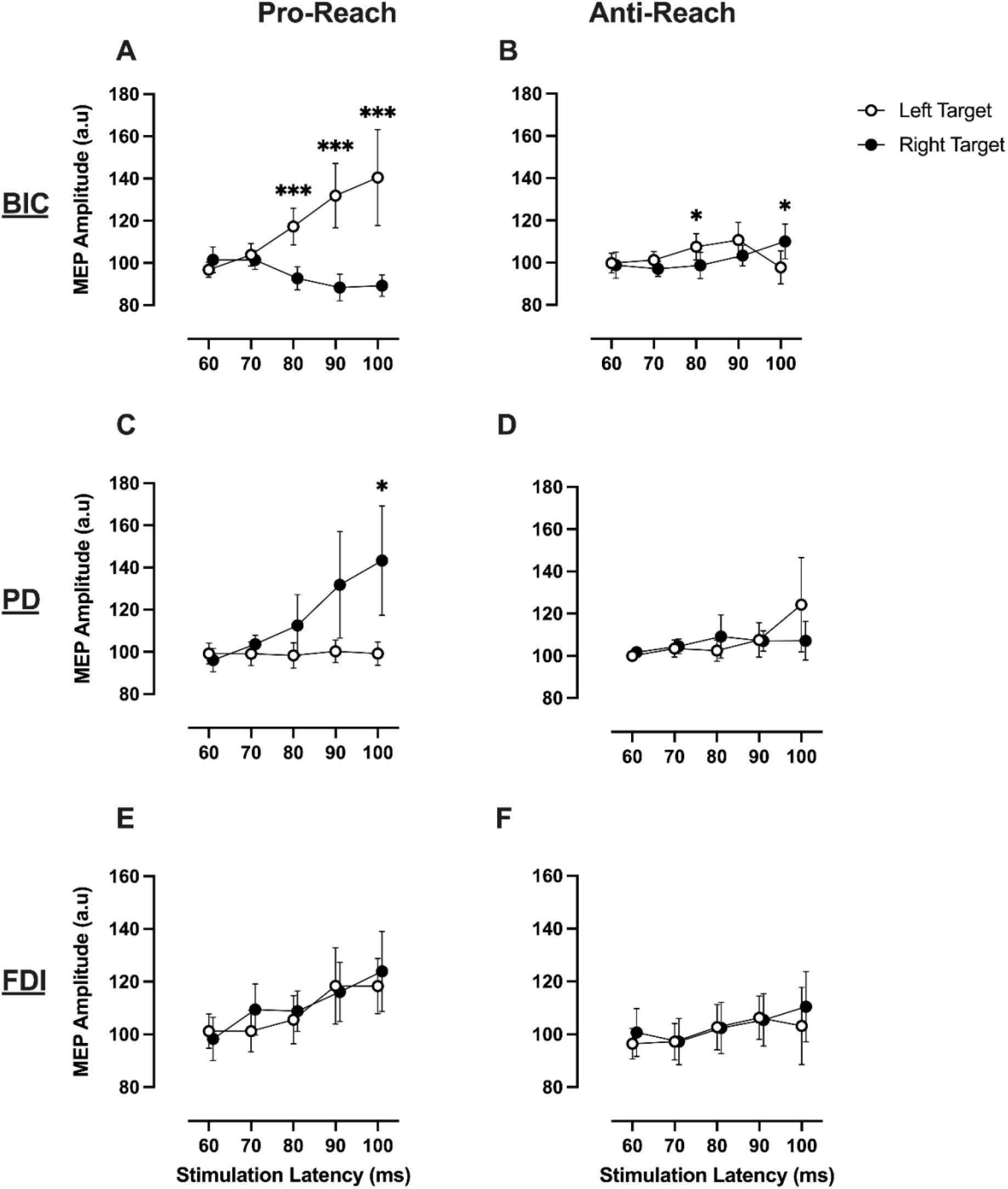
Time course of MEP amplitude as a function of stimulation latency for Posterior Deltoid (***A&B***), Biceps Brachii (***C&D***), First Dorsal Interosseous (***E&F***). First column shows results in the pro-reach condition and second column shows results in the anti-reach condition. Black circles represent the left target direction and white circles represent the right target direction. Error bars show 95% CI. *P < 0.05, **P < 0.01, ***P < 0.001.

The PD muscle MEPs showed a mirror-opposite trend to that of its antagonist PEC muscle in both pro- and anti-reach conditions (**Figure 6C&D**). In the pro-reach condition, the MEP amplitude was dependent on the location of the target and the stimulation latency (main effect of target direction: *F*_1,19_ = 5.7, *P* = 0.028; main effect of stimulation latency: *F*_1.5,27.6_ = 8.6, *P* = 0.003; interaction: *F*_1.5,28.3_ = 7.6, *P* = 0.005). At 100 ms, the average MEP for right target was larger than the left target (*P* = 0.032); this is expected as PD is the agonist for right reaches. In the anti-reach condition, the MEP amplitude showed a main effect of stimulation latency (*F*_1.4,26.7_ = 6.4, *P* = 0.011). However, there was no significant effect of the target direction (*F*_1,19_ = 0.4, *P* = 0.553) or target direction by stimulation latency interaction (*F*_1.4,27.2_ = 1.9, *P* = 0.173). Despite this non-significant effect, **Figure 6D** shows a pattern in PD that is qualitatively similar to that observed for PEC, with the MEP amplitudes at 80 ms apparently biased towards the direction of the target and flipping in the opposite direction at 100 ms. It is important to note here that the PD muscle wasn’t preloaded in this study, which might have contributed to the non-significant target-oriented responses at 80 ms.

By contrast, MEPs in the FDI muscle in the hand were not significantly changed by target or reach direction in either pro or anti-reach conditions (**Figure 6E&F)**. In pro-reach conditions, there was no main effect of target direction (*F*_1,19_ = 0.9, *P* = 0.364) or target direction by stimulation latency interaction (*F*_2.8,52.9_ = 0.9, *P* = 0.424), however, there was an effect of stimulation latency (*F*_2.5,46.9_ = 10.3, *P* < 0.001). This could be a general increase in excitability of the corticospinal tract that is not directionally specific as the participant prepares to move. In the anti-reach condition, there was no main effect of the target direction (*F*_1,19_ = 1.0, *P* = 0.327), stimulation latency (*F*_1.9,37.7_ = 1.9, *P* = 0.157), or target direction by stimulation latency interaction (*F*_3.2,60.6_ = 0.5, *P* = 0.674). The data show that muscles that are functionally relevant to the reaching task (PEC, PD and BIC) show early target-oriented corticospinal excitability changes, whereas the excitability of projections to hand muscles that are not functionally relevant for the reach (FDI) are unaffected by the target direction.

### *Experiment 2 –* No significant differences between TMS and TES probes of early corticospinal excitability

In experiment two, we sought to establish whether the early target-oriented changes in corticospinal excitability seen in experiment one were driven by cortical or subcortical modulation. To this end, we stimulated the motor cortex using either transcranial magnetic stimulation (TMS) or transcranial electrical stimulation (TES) at 60, 80, or 100 ms after the onset of the visual target. Stimulation was also given just prior to the onset of the reach; this stimulation latency was called the *voluntary* latency. We tracked the change in MEP as a function of stimulation latency for the two reach types (pro- and anti-reach) and two target directions (left and right). If the target directed changes are mediated within the motor cortex, there should be a dissociation between the target-related modulation of TMS and TES responses. On the contrary, if modulation occurs sub-cortically, there should be no differences in the timing or magnitude of target-directed MEP modulations.

First, we examined how participants performed in experiment two. Participants generally made rapid reaches to the nominated direction in both pro and anti-reach conditions (reaction time: pro-reach = 199 ± 25 ms, anti-reach = 232 ± 28 ms). Reaction times were significantly slower when the reaches had to be made away from the target location (main effect of reach-type: *F*_1,19_ = 52.6, *P* < 0.001; **Figure 7A**). Although the main effect of target direction was non-significant (*F*_1,19_ = 0.02, *P* = 0.885), the significant interaction between target direction and reach type (*F*_1,19_ = 12.2, *P* = 0.002) shows that the target direction also had an effect on participant’s reaction time in the pro- and anti-reach conditions (**Figure 7A)**. Reaction time was also significantly affected by brain stimulation (RM one-way ANOVA: *F*_1.6,30.1_ = 9.5, *P* = 0.001). Participants initiated movement earlier on TES trials (211 ± 23 ms) compared to *noStim* (226 ± 26 ms, *P* = 0.003) and TMS trials (219 ± 27 ms, *P* = 0.006) (**Figure 7B**). This is consistent with a previous study which showed that electrical stimulation can induce the early release of motor actions (Marinovic et al., 2015). The occurrence of reach errors was not significantly different between pro- and anti-reaches (*t*_19_ = 2.0, *P* = 0.058; **Figure 7B)**, or affected by target direction (*t*_19_ = 1.7, *P* = 0.106). Participants also made more errors on trials with brain stimulation compared to trials without brain stimulation (RM one-way ANOVA: *F*_1.7,31.9_ = 34.5, *P* < 0.001; **Figure 7D**).

**Figure 7.**
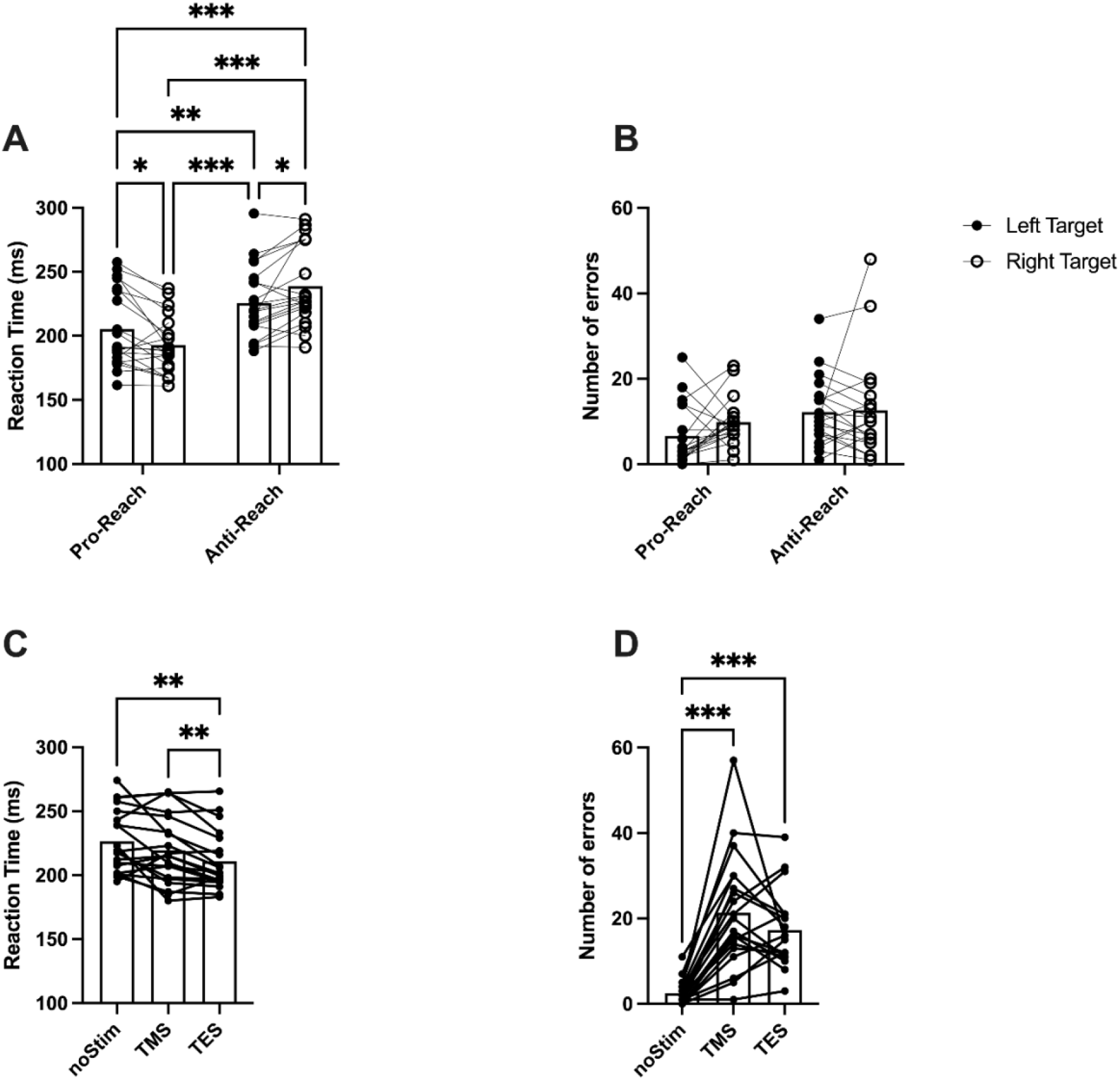
***A***: Reaction time of correct reaches in the pro-reach and anti-reach condition. ***B***: Reaction time in trials with and without brain stimulation. ***C***: Number of reach errors in pro- and anti-reach condition ***D***: Number of reach errors in trials with and without brain stimulation. Error bars show 95% CI. *P < 0.05, **P < 0.01, ***P < 0.001.

The primary objective of this experiment was to establish whether the size of MEPs generated by TMS and TES was differentially modulated within the first 100 ms after presentation of a visual target, which would imply a cortical contribution to express visuomotor responses. **Figure 8** shows example MEP traces from an exemplar participant in the pro-reach condition, using both TMS and TES. The average size of MEP evoked by TMS (38.7 ± 16.9 μV) was not significantly different from the TES (42.6 ± 16.2 μV) (*t*_19_ = 1.8, *P* = 0.086). On average the latency of the MEPs evoked by TES (6.91 ± 0.92 ms) was significantly shorter than TMS (8.01 ± 1.07 ms) (*t*_19_ = 4.03, *P* < 0.001). This indicates that the two forms of stimulation activated the corticospinal tract at distinct sites, and is consistent with previous reports of an additional synaptic delay for TMS (Day et al., 1989; Di Lazzaro et al., 1998a). For this representative subject, the time course of MEP modulation appeared to be similar for TMS and TES. We observed a clear increase in MEP amplitude for the left target as stimulation was delivered at later times, while MEP amplitude decreased for the right target. This finding is consistent with our results from experiment one.

**Figure 8.**
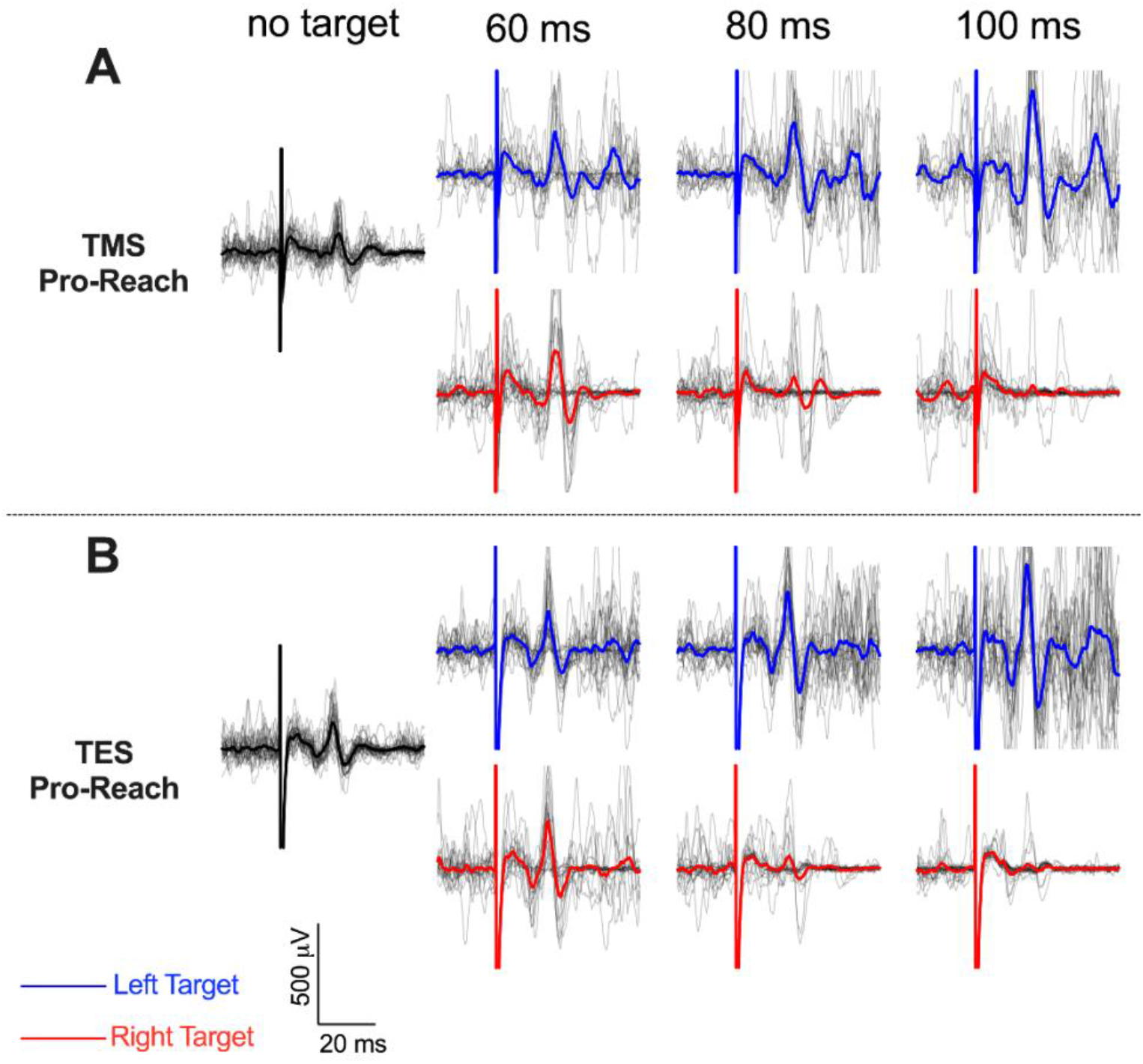
Individual MEP traces from an exemplar participant in the pro-reach condition with the average MEP waveform overlaid on top of it. **A**: Top two rows show traces from the TMS condition. **B**: bottom two rows show traces from the TES condition. MEP recorded from the pectoralis major muscle.

To test for target-related modulation of MEP size between TES and TMS evoked responses, we quantified how the difference in mean amplitude between the left and right MEPs varied with stimulation latency and stimulation type. This difference score is essentially a measure of the degree to which responses to brain stimulation were modulated by the location of the target. A difference score value above zero would indicate that a left-side target produced larger MEPs than a right-side target and a value below zero would indicate larger MEPs for the right-side target. We ran a RM-ANOVA on the difference scores with stimulation latency (60, 80 and 100 ms) and stimulation type (TMS and TES) as the two factors. This analysis was performed separately on all four muscles and for the pro- and anti-reach tasks. If MEPs are modulated within motor cortex according to target location, there should be significant interaction effect between stimulation type and stimulation latency.

For the pectoralis major (PEC) and biceps brachii (BIC) muscle, we observed difference score patterns for both types of stimulation that are consistent with the target-oriented response seen in experiment one (**Figure 9A-D)**. In both pro- and anti-reach conditions, TMS and TES resulted in positive difference scores at 80 ms indicating the presence of target-biased excitability changes. At 100 ms, consistent with what we observed in experiment one, the score remained positive for pro-reaches but flipped to a negative value for the anti-reaches. MEPs were modulated based on the latency of stimulation as evidenced by a significant main effect of stimulation latency in both pro- (PEC: *F*_1.3,25.4_ = 21.5, *P* < 0.001; BIC: *F*_1.2,22.3_ = 15.2, *P* < 0.001) and anti-reach (PEC: *F*_1.3,24.3_ = 15.5, *P* < 0.001; BIC: *F*_1.6,29.9_ = 8.9, *P* = 0.002) conditions. For both PEC and BIC muscles, however, there were no significant differences in the way that MEPs elicited by TMS and TES were modulated across pro- and anti-reach conditions. There was no significant main effect of stimulation type, nor an interaction effect between stimulation type and timing for the pro reach condition (main effect: *F*_1,19_ = 1.0, *P* = 0.339; interaction: *F*_1.9,36.1_ = 0.6, *P* = 0.541) or the anti-reach condition (main effect of stimulation type: *F*_1,19_ = 2.4, *P* = 0.136; interaction effects: *F*_1.7,31.7_ = 0.1, *P* = 0.896) for the PEC muscle. The BIC muscle similarly showed was no significant main effect of stimulation type nor an interaction effects in pro- (main effect of stimulation type: *F*_1,19_ = 3.4, *P* = 0.08; interaction: *F*_1.4,27.1_ = 0.2, *P* = 0.746) or anti-reach condition (main effect of stimulation type: *F*_1,19_ = 1.7, *P* = 0.211; interaction effects: *F*_1.7,32.2_ = 2.3, *P* = 0.128).

**Figure 9.**
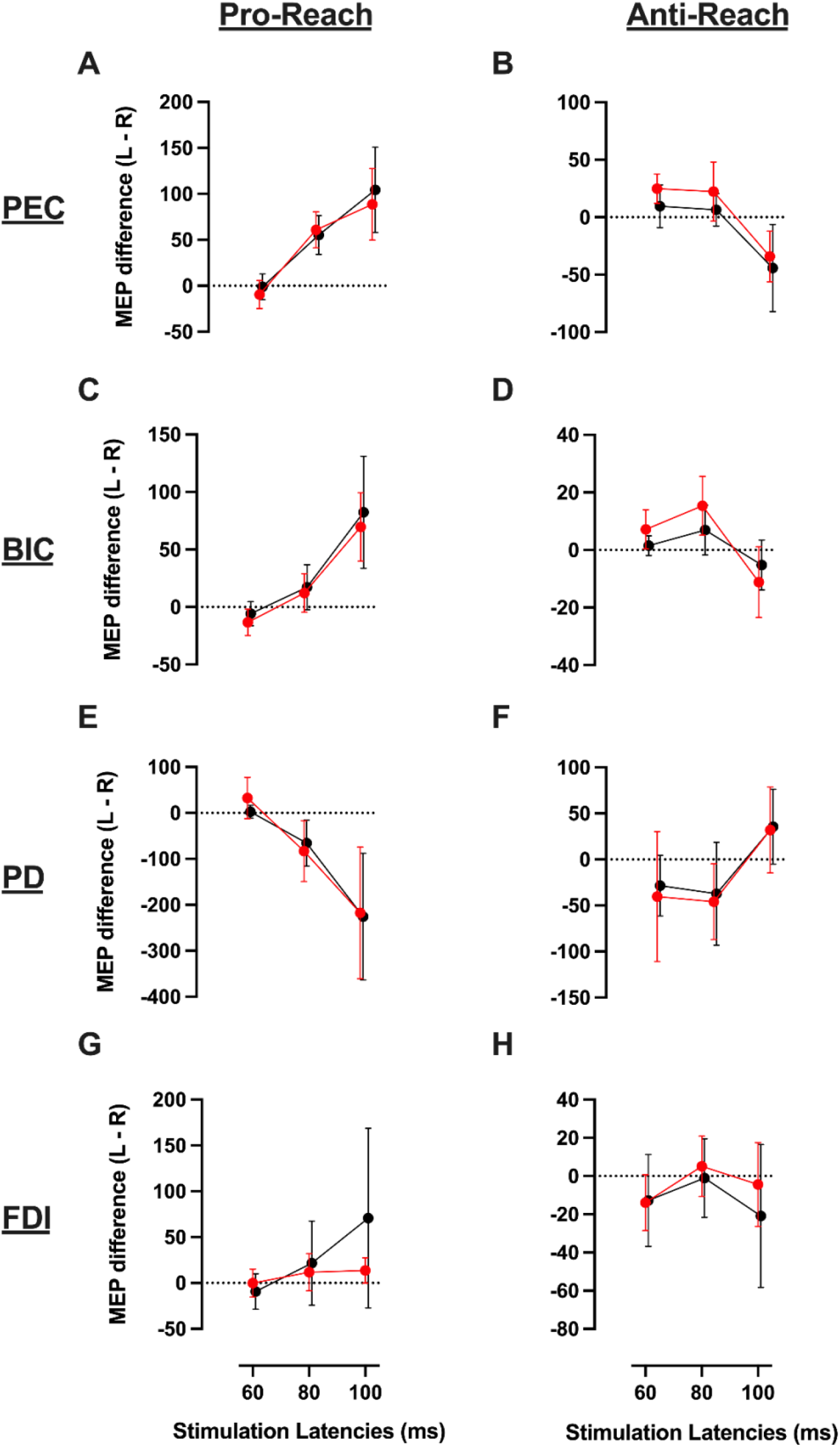
***A***: Time course of change in MEP amplitude difference between left and right targets (L-R) as a function of stimulation latency for Pectoralis Major (***A&B***), Posterior Deltoid (***C&D***), Biceps Brachii (***E&F***), First Dorsal Interosseous (***G&H***). First column shows results in the pro-reach condition and second column shows results in the anti-reach condition. Black circles represent Transcranial Electrical Stimulation (TES) and red circles represent the Transcranial Magnetic Stimulation (TMS).

In the antagonist posterior deltoid (PD) muscle (**Figure 9E&F**), we observed an opposite pattern to that of the PEC and BIC muscles. In both pro- and anti-reach conditions, TMS and TES resulted in negative difference scores at 80 ms, demonstrating muscle activity reciprocal to that seen in the PEC and BIC. There were also no significant differences in the way that MEPs elicited by TMS and TES were modulated across pro- and anti-reach conditions. There was a significant main effect of stimulation latency in the pro-reach condition (*F*_1.0,19.5_ = 11.9, *P* = 0.002) but not in the anti-reach condition (*F*_1.1,21.0_ = 3.7, *P* = 0.066), but neither the pro- or anti-reach conditions had a significant main effect of stimulation type (Pro: *F*_1,19_ = 0.1, *P* = 0.741, Anti: *F*_1,19_ = 0.8, *P* = 0.394) or a significant interaction effect (Pro: *F*_1.5,28.0_ = 0.7, *P* = 0.476, Anti: *F*_1.3,24.5_ = 0.04, *P* = 0.890).

For the first dorsal interosseous (FDI) muscle (**Figure 9G&H**), there was no significant main effect of stimulation latency (Pro: *F*_1.2,23.5_ = 2.6, *P* = 0.113, Anti: *F*_1.7,32.9_ = 2.5, *P* = 0.108), stimulation type (Pro: *F*_1,19_ = 1.3, *P* = 0.268, Anti: *F*_1,19_ = 0.3, *P* = 0.596) or interaction effects (Pro: *F*_1.1,21.1_ = 1.4, *P* = 0.260, Anti: *F*_1.6,29.5_ = 0.5, *P* = 0.550).

In summary, arm muscles that are functionally relevant to executing the reach showed difference-score patterns that are consistent with the presence of target-oriented excitability changes, whereas the excitability of projections to a functionally irrelevant hand muscle was not modulated. This observation is consistent with the results of experiment one. We also saw that there were no significant differences in MEP modulation between TMS and TES across stimulation latencies in any functionally relevant muscle. This suggests that the early target-oriented corticospinal excitability changes likely occurred downstream of the corticospinal output cells, supporting the idea that subcortical structures are responsible for express visuomotor responses.

In experiment one, when we measured target-directed excitability changes every 10 ms between 60 and 100 ms after target presentation, we saw that the earliest target-driven excitability changes in the biceps muscle occurred at 80 ms (**Figure 6A-B**), compared to 70 ms for the pectoralis muscle (**Figure 5A-B**). Therefore, it was crucial to compare TMS and TES responses at 80 ms after target presentation for the biceps muscle in experiment 2, at the earliest latency of excitability change. If the earliest increase in excitability were driven by increased cortical excitability, we would have expected an earlier increase in responsiveness to TMS than to TES. Thus, the lack of difference between TMS and TES at 80 ms for the biceps muscle provides additional evidence that target-directed corticospinal excitability changes are driven by subcortical modulation (**Figure 9C-D**).

## DISCUSSION

The key findings were that the earliest changes in corticospinal excitability in a visually guided reaching task reflect a bias to move the limb towards the physical location of the target regardless of task instructions, and that these excitability changes were likely mediated subcortically given the similar modulation patterns for TMS and TES responses. Excitability changes were evident as early as 70 ms after target presentation, and appear to reflect a rapid visuomotor transformation process that generates a target-oriented motor output rather than a goal-oriented motor output. The absence of target-specific modulation of corticospinal responses in a hand muscle suggests that early target-oriented visuomotor processing is selective to functionally relevant muscles, however, it is also possible that the neural structures responsible for such early visuomotor transformation do not project to distal muscles in general.

An interesting finding was that the tuning of corticospinal excitability in the anti-reach condition flipped from being target-oriented at ∼80 ms after target presentation to being goal oriented at ∼100 ms after target presentation. This was observed across multiple muscles in both experiments (**Figure 5B**, **Figure 6B&D**, **Figure 9B&D&F**). It is tempting to interpret this reorientation in excitability as evidence for rapid visuomotor transformation that reflects the anti-reach task goal. In this case, the fact that the flip occurred for both TMS and TES responses would suggest that subcortical circuits are capable of implementing a non-veridical task rule to reach away from a physical target. However, there are alternative explanations that might apply. For example, in a delayed reaching task, Wood et al. 2015 found that target-directed express visuomotor responses in the pectoralis major muscle were immediately followed by a “rebound” muscle activity in the opposite direction. Such “rebound” effects are to be expected when there is a brief and transient activation or inhibition of a pool of tonically firing motoneurons, due to synchronization of motoneuronal membrane potential trajectories. It is also possible that oscillating responses could reflect “low-level” dynamics intrinsic to the visuomotor circuits that generate express behaviour, potentially due to reciprocal inhibition between neural populations representing alternative target locations. Thus, further investigation is needed to understand the characteristics and relevance of the rapid reorientation of corticospinal excitability observed here, and the time-course of true “movement-directed” corticospinal processing in anti-reach tasks.

The visual target tuning of corticospinal excitability changes reported here is consistent with recent studies that reported target-oriented express visuomotor responses in the 80-120 ms window after the onset of a visual target (Corneil et al., 2004; Wood et al., 2015; Gu et al., 2016, 2018, 2019; Atsma et al., 2018; Glover and Baker, 2019; Kozak et al., 2019; Contemori et al., 2020, 2021, 2022b; Kearsley et al., 2022). Similar to the excitability changes seen in our study, these express visuomotor responses are also tuned to the direction of the visual target and not the direction of the reach (Gu et al., 2016). However, we did not detect consistent express visuomotor responses in our participants as defined by standard criteria. Despite the absence of robust express visuomotor responses, we found some target-related EMG changes in pectoralis muscle within the express response time-window (**Figure 2**). Although this implies that the observed MEP modulations were influenced by changes in motoneuron activation, the time-course of target-directed modulation is better defined for MEPs than background EMG (**Figure 5A**), and there are some discrepancies between the time courses of EMG and MEP modulations. This suggests that TMS-induced MEPs were subject to influence by non-motoneuronal aspects of corticospinal processing during the early stages of reaching.

The timing of corticospinal excitability effects induced by visual stimulation observed here is broadly consistent with previous reports by Nakajima et al. 2021 and Suzuki et al. 2021. Nakajima et al. 2021 found that a bright visual flash presented centrally when the arm was stationary and under tonic contraction increased the size of MEPs elicited in the biceps brachii muscle by both TMS and TES from 60 ms (Nakajima et al., 2021). Suzuki et al. 2021 used a target jump paradigm during an ongoing reach and found modulation in MEP size from 70 ms in the muscles involved in making reach corrections that depended on the direction of the target jump (Suzuki et al., 2021). Several findings suggest that these rapid modulations of corticospinal responsiveness were likely mediated subcortically. Firstly, both studies found similar evoked response modulations using TMS, TES and cervicomedullary stimulation. Secondly, Suzuki and colleagues found that MEPs in the triceps muscle increased at 70 and 80 ms in a person with hemianopsia, even when the target jumped to their “blind” right hemifield. Thirdly, both studies found that MEPs to combined TMS and ulnar nerve stimulation were also enhanced in the presence of visual stimulation, which suggests a visually driven modulation of cervical propriospinal interneuron states. Based on this evidence, the authors suggested that these flexible MEP modulations were due to a short-latency pathway that bypasses the primary visual cortex and runs directly from the retina to the cervical spinal interneurons via a subcortical pathway. The data from our study are consistent with this interpretation. However, fast visual cortex to superior colliculus pathways are known to exist in monkeys (Kadoya et al., 1971; Schiller et al., 1974; Fries, 1984; Cusick, 1988; Lock et al., 2003; Collins et al., 2005; Baldwin and Kaas, 2012; Cerkevich et al., 2014), and the potential for an analogous pathway to contribute to early visuomotor excitability changes in humans cannot be ruled out. Regardless, the consistent finding that the time course of early MEP modulations caused by a visual target does not differ between TMS and TES, indicates that evoked muscle responses to corticospinal tract stimulation are similar irrespective of whether stimuli are subject to modulation at cortical or subcortical levels. Thus, the excitability changes caused by the visual target likely occur downstream of the corticospinal output neurons. Taken together, these studies strongly indicate an express subcortical pathway that bypasses the motor cortex and initiates movements towards visual targets.

A prime candidate for such a subcortical pathway involves the superior colliculus and the reticulospinal tract. The superior colliculus is a midbrain structure that receives direct projections from the retina and can encode the spatial location of a salient visual stimulus (Basso and May, 2017). Output signals from the superior colliculus are known to project to the reticular formation to produce express motor responses, such as in the case of express saccades (Moschovakis et al., 1996; Munoz et al., 2000). Although the dynamics of limb control are more complicated than those of eye movements, there is evidence that the superior colliculus and the reticulospinal tract contribute to reaching behaviour in non-human animals. The activity of reach-related neurons in the superior colliculus and the reticular formation is modulated during reaching (Werner, 1993; Kutz et al., 1997; Werner et al., 1997a, 1997b) and associated muscle activation (Werner et al., 1997a; Stuphorn et al., 1999). Electrically stimulating neurons in the deep layers of the superior colliculus can lead to short-latency changes in the limb movement trajectories (Philipp and Hoffmann, 2014). Converging evidence also suggests that the reticulospinal tract, which is a downstream target of superior colliculus, is crucial for executing reaching behaviour. Lesion studies in cats and monkeys show that animals with an intact reticulospinal tract can recover and produce goal-directed reaching behaviour, when other major pathways (i.e. corticospinal and rubrospinal) to the muscle are lesioned (Alstermark and Isa, 2012; Alstermark and Pettersson, 2014). Studies in mice and monkeys that implemented viral and optogenetics tools also showed that the brainstem and its downstream targets are crucial for reaching and grasping behaviour (Kinoshita et al., 2012; Azim et al., 2014b; Esposito et al., 2014; Ruder et al., 2021). The extensive connectivity between the brainstem and cerebellum means that ongoing movement can be refined using feedback and learning (Azim et al., 2014a; Alstermark and Ekerot, 2015; Azim and Alstermark, 2015). Furthermore, mathematical modelling of spinal interneuron networks have shown that the spinal circuitry is capable of the computations required to account for limb movement dynamics across a wide range of tasks (Raphael et al., 2010; Tsianos et al., 2014). Moreover, recent experiments show that even the simplest spinal reflexes can be functionally modulated by posture (Weiler et al., 2019). Thus, the results of the present study add to a growing body of evidence consistent with the idea that primitive subcortical pathways play an important role in the production of reaching behaviour.

### Conclusions

The results show that the responsiveness of corticospinal projections to proximal limb muscles invariably reflects the visual target rather than the reach direction within 70ms of target presentation. The similarity in time-course and magnitude of response modulations to TMS and TES suggests that these early target-directed changes in MEP size occur at a spinal rather than cortical level of the neuroaxis. A subcortical pathway involving the superior colliculus and reticulospinal tract is a prime candidate for generating the target-oriented signals that cause these early excitability changes.

## METHODS

### Participants

Forty participants without a history of neurological disorder participated in this study. Twenty participants (11 males, 9 females; mean age: 27.2, SD: 7.3; 16 right-handed, 4 left-handed) participated in the first experiment and twenty participants (10 males, 10 females; mean age: 26.3, SD: 6.2; 17 right-handed, 3 left-handed) participated in the second experiment. Four participants who participated in the first experiment also took part in the second experiment. All participants provided their informed consent after they read an information document and received a verbal explanation, and filled out a health screening questionnaire to make sure they were not contraindicated for TMS/TES. They were free to withdraw from the experiment at any time. The experimental procedures conform to the Declaration of Helsinki, and the University of Queensland Medical Research Ethics Committee (Brisbane, Australia) approved the procedures.

### General setup

Participants were seated in front of a horizontal desk surface, with their forearm resting on a custom-built air sled which minimized friction during reaching (**Figure 10A**). The chair height was adjusted to allow reaching movements in the transverse plane, from a starting position of ∼90° shoulder flexion. Participants were instructed to initiate the reaches from the shoulder joint, instead of bending their elbows. A custom-made pulley mechanism attached to the air-sled applied a load force of ∼5N (towards right) on the right arm to increase the baseline activity of the pectoralis major muscle. Participants rested their head in a video-based eye tracking system mounted on the experiment table (EyeLink 1000, SR Research Ltd., Ontario, Canada), this offered chin and forehead support to ensure head stability during reaching movements. Eye tracking data (sampled at 1000 Hz) from the EyeLink system were used to initiate a trial by detecting eye fixation. Participants viewed the visual stimuli on an LCD monitor (BenQ XL2720, refresh rate 120 Hz) that was calibrated to ensure a linear increase in luminance across the greyscale. The monitor was positioned ∼57 cm in front of the eye, so that 1 cm on the screen equaled ∼1-degree visual angle (dva). Once the participants were comfortably seated, surface Electromyographic (EMG) electrodes (24 mm disposable Ag-AgCl electrodes) were attached to the clavicular head of the pectoralis major muscle, biceps brachii muscle, posterior head of the deltoid muscle, and first dorsal interosseous of the right arm. The EMG signals were amplified (x1000) and band-pass filtered (0.1-10 kHz) using Grass P511 isolated amplifiers. A three-axis accelerometer (Dytran Instruments, Chatsworth, CA) taped to the right index finger measured the acceleration of the arm. EMG and accelerometer data were digitized at 5000 samples/s and stored on a computer using a 16-bit analog-digital converter (USB-6343-BNC DAQ device, Accelerometer data was analyzed to detect the onset and direction of the arm movement.

**Figure 10.**
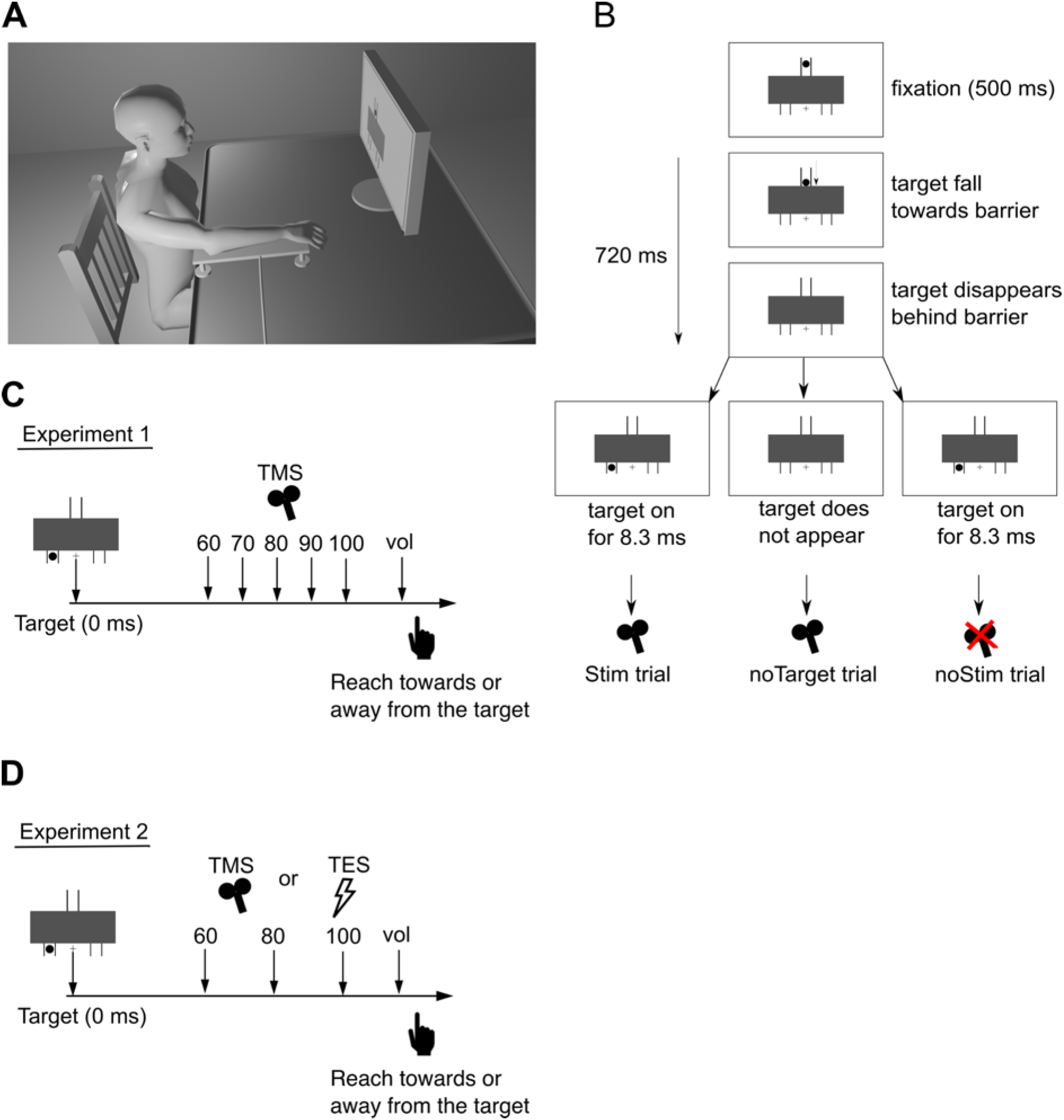
***A:*** Experiment setup. ***B:*** Emerging target paradigm used in both experiments. ***C:*** In the first experiment, transcranial magnetic stimulation (TMS) was delivered to the left motor cortex at either 60, 70, 80, 90, 100 or *voluntary* latency (30 milliseconds prior to the median onset latency of the voluntary muscle activity). ***D:*** In the second experiment, either transcranial magnetic stimulation (TMS) or transcranial electrical stimulation (TES) was delivered to the left motor cortex at either 60, 80, 100 or a *voluntary* latency (30 milliseconds prior to the median onset latency of the voluntary muscle activity).

National Instruments, Austin, TX).

In both experiments 1 and 2, we used an emerging target paradigm (**Figure 10B**) that has been shown to promote short latency muscle responses (Contemori et al., 2020; Kozak et al., 2020). In this paradigm, a visual target which was a black filled circle (diameter: ∼ 2 dva; luminance: ∼0.3 cd/m^2^) moved down an inverted y-shaped track (color: black) set against a gray background (∼170.4 cd/m^2^). Participants fixated on a cross (∼1 dva; color: black) situated midway between the left and right y-track. A part of the path was occluded by a rectangular barrier. From a starting position ∼ 5 dva above the barrier, the black circle dropped at a constant velocity of ∼ 35 dva/s towards the barrier and gradually disappeared behind it. It then reappeared as a whole circle beneath the barrier on either the left or right “arm” of the y-track (fixation spot to target distance: ∼10 dva) for one screen refresh cycle (∼8.3 ms); this gives the appearance of a flashing target. Note that the target did not reappear gradually; the entirety of the target flashed when it reappeared under the barrier. Previous work showed that transient targets elicit stronger express visuomotor responses than continuously moving targets (Contemori et al., 2020). The target took ∼ 720 ms to travel from its start position at the top of the track to its final position just below the barrier. This paradigm was slightly different in the two experiments (see below).

To start a trial, participants brought their arm to the centre of the workspace to point at the fixation and held their gaze on the fixation cross. Once the eye tracker detected a 500 ms long unbroken fixation, the black target moved towards the barrier at a constant velocity and disappeared behind it, before flashing underneath the barrier, either on the left or right arm of the y-track. If participants broke fixation before the target appeared, an error message was shown and the trial was repeated. Target direction was randomized within each reaching block. Maintaining a constant velocity for the target ensured that the time of appearance of the target beneath the barrier was predictable on every trial. This temporal predictability has been shown to facilitate the short latency muscle response (Contemori et al., 2020). Once participants saw the target, they executed a fast reach either towards the target (in pro-reach trials) or away from the target (in anti-reach trials). Pro- and anti-reach trials were performed in separate blocks to simplify the task, as increasing task complexity can diminish express visuomotor responses (Gu et al., 2018).

### Brain Stimulation

Transcranial Magnetic Stimulation (TMS) was delivered using a figure of eight coil (wing diameter of 8 cm) connected to a MagStim 200^2^ stimulator (The MagStim Company, Carmartheshire, Wales, UK). Transcranial Electrical Stimulation (TES) was delivered using a Digitimer D180A stimulator (Digitimer, L. Welwyn Garden City, UK). To find the optimal location on the scalp that produced consistent motor evoked potentials (MEPs) from the pectoralis major muscle, we first measured the participant’s head to locate the Cz electrode location (EEG 10-20 system). With the center of the TMS coil starting from a spot 2 cm anterior and 3 cm lateral to the Cz site, single pulses were delivered to the scalp. We adjusted the coil position and stimulator output intensity until consistent MEPs were induced in the pectoralis major muscle. The coil was oriented anterior-posteriorly, approximately 45° with respect to the sagittal plane and held tangential to the scalp. Note that the pectoralis muscle was loaded using a pulley and weight system, which increased the baseline activity of the muscle and lowered the stimulation intensity required to induce a MEP (Di Lazzaro et al., 1998b).

The following steps were performed only for the second experiment. Once the optimal scalp spot was found, TES electrodes were placed on the scalp with cathode on the Cz location, and anode on the previously found optimal spot. Prior to placement, electrode cream (EC2 Electrode Cream, Grass Instrument Company) was applied to 9mm Na/NaCl electrodes to reduce impedance. The width of the electric pulse was either 50 µs or 100 µs depending on participant tolerance and response efficacy. The stimulation intensity was subsequently adjusted to produce consistent MEPs from the pectoralis major muscle of comparable magnitude to the TMS responses.

An Arduino Uno development board in conjunction with a photodiode was used to ensure precise timing of the stimulation delivery. The photodiode sent a pulse to the Arduino board when it detected a visual target. The Arduino board then triggered the TMS or TES device at the stimulation latency provided to it by the experiment computer at the start of every trial (1ms precision).

#### Experiment 1

In experiment one, we aimed to test how long after target presentation the corticospinal excitability was modulated according to the target location with respect to the starting hand position. We systematically stimulated the motor cortex using transcranial magnetic stimulation (TMS) at a range of latencies from target presentation, as participants performed either pro-reaches (reach towards the target) or anti-reaches (reach opposite of the target). Experiment one consisted of three sessions completed on the same day.

To start session one, participants performed 4 *baseline* reaching blocks without receiving any brain stimulation. There were two blocks of pro-reaches and two blocks of anti-reaches. We collected 50 trials each for the four conditions tested: 2 target locations (left or right) * 2 reach types (pro- or anti-reach), for a total of 200 trials. After the *baseline* blocks, we used transcranial magnetic stimulation (TMS) to identify the scalp location for producing consistent motor evoked potentials (MEP) from the pectoralis major muscle (see Methods: **Brain stimulation** for more details). This concluded session one.

Sessions two and three each contained seven blocks. The first block in both sessions involved TMS with no targets or movement. There were 20 trials in these blocks, thus over two sessions 40 trials were recorded. The purpose of TMS blocks with no movement was to measure the baseline MEP in the pectoralis major muscle when participants were not reaching and just holding a steady background contraction with the y-track, barrier and fixation cross shown on screen. Participants were instructed that no target would be shown to them, and they just had to hold their arm pointing at the fixation cross. Note that the pectoralis major muscle is activated by doing this, as it is working against the force exerted by the pulley on the arm. As participants held this arm position, 20 TMS pulses were delivered to the marked scalp location at an inter-pulse interval of ∼ 5 sec (0.5 – 1 second jitter) to record the MEPs.

The remaining six blocks in each session involved TMS during reaching. There were 68 trials in each of these blocks, thus over two sessions 816 trials (12 blocks * 68 trials) were recorded. Participants were given on average 30 minutes (minimum 15 minutes and maximum 1 hour) break between sessions two and three to rest. In the TMS blocks with reaching, participants performed fast reaches as they received brain stimulation. All blocks in each session were made up either entirely of pro-reaches or entirely of anti-reaches. This was randomly assigned and counterbalanced across participants. Each block contained three different types of trial (**Figure 10B**): 1) *Stim* trials in which TMS pulses were delivered at various time points after target presentation; 2) *noStim* trials in which TMS pulses were not delivered after target presentation; 3) *noTarget* trials in which TMS pulses were delivered, but the target did not reappear beneath the barrier after disappearing behind it. Participants were instructed to withhold the reach in *noTarget* trials. The purpose of the *Stim* trials was to track when corticospinal excitability reflected the target identity and when it reflected the reach direction. TMS pulses were delivered at six latencies after visual target presentation in the pro- and anti-blocks; five of which were common for both reach types across all participants: 60, 70, 80, 90 and 100 ms (**Figure 10C**). The sixth latency known as the “voluntary” latency was given just prior to the onset of the reach and was intended to record the MEPs when the excitability of the cortex was expected to be highly modulated by the task goal. The sixth latencies for pro- and anti-reaches were calculated based on the per-participant median EMG onset latency of pectoralis muscle for left-target pro-reach and right-target anti-reach respectively. The median onset latency was calculated in session one using the EMG data from the *baseline* blocks. The EMG onset latency was defined as the time when the EMG signal deviated more than five standard deviations from the mean EMG in the baseline period (100 ms window prior to target presentation). We picked two median onset latencies: 1) when left reaches were executed towards left targets (pro-reach) and 2) when left reaches were executed towards right targets (anti-reach). We subtracted 30 ms from these latencies to obtain the two stimulation timings for pro- and anti-reaches, which we termed as the *voluntary* latencies. Thus, the pro- and anti-reach *voluntary* latencies were determined from median EMG onset latency from the left reaches and applied for TMS timing during both left and right reaches. Using this method, the median *voluntary* stimulation latency for pro-reach was 145 ms (range: 110-220); and for anti-reach was 170 ms (range: 130-240).

In the *noStim* trials, participants were not given the TMS pulse after target presentation. The purpose of these trials was to measure muscle activity during the reaching task in the expectation of stimulation, but without stimulation being applied. In the *noTarget* trials, stimulation was delivered 800 ms after the target started dropping towards the barrier, or 80 ms after the expected “target onset” had the target appeared beneath the barrier. In this way, we could investigate how the TMS pulse influenced the corticospinal tract when the participant was prepared to move but received no target information. Participants were unaware when the *noTarget* trials would occur and were instructed to withhold the reach if no target appeared under the barrier. The purpose of these trials was to understand the contribution of visual target information to corticospinal excitability, as opposed to potential excitability changes caused by target expectation and motor preparation. To summarize, the *Stim* trials had 24 conditions: 6 latencies (60, 70, 80, 90, 100, voluntary) x 2 (left and right target) x 2 (pro and anti-reach); *noStim* trials had 4 conditions: 2 (left and right target) x 2 (pro and anti-reach); and *noTarget* trials had 2 conditions: pro and anti-reach. Every *Stim* and *noStim* condition had 27 trials each, and the two *noTarget* condition had 30 trials each. Therefore, 648 *Stim* trials, 108 *noStim* trials and 60 *noTarget* trials were distributed across the 12 stimulation blocks.

#### Experiment 2

In experiment two, we tested whether the pattern of MEP modulation at latencies within 100 ms of target presentation differed significantly between TMS and TES. A significant difference between TMS and TES response modulation would suggest that the early target-oriented corticospinal excitability changes were at least partly driven by modulation within the motor cortex. To test this, we probed the responsiveness of the corticospinal pathway close to the express response window and around the onset of voluntary EMG with both TMS and TES. The general timeline of experiments one and two were similar, in that both involved three sessions. However, in experiment 2, participants performed sessions one and two on the same day, but session three was done on a different day. In the first session, participants performed four *baseline* blocks of reaching trials prior to any brain stimulation. We collected 50 trials each for the four conditions tested: 2 target locations (left or right) * 2 reach types (pro- or anti-reach), for a total of 200 trials. After this the ideal scalp location for stimulation was identified (see Methods: **Brain stimulation** for more details). Both the second and third sessions had one stimulation block with no movement and six stimulation blocks involving reaching. There were 40 trials in each no movement block comprising of 20 TES and 20 TMS trials in pseudorandomized order. In the six reach stimulation blocks in each session, participants received randomized *Stim* and *noStim* trials. All blocks in each session were made up either entirely of pro-reaches or entirely of anti-reaches. The task order was randomly assigned and counterbalanced across participants. There were 72 trials in each of these blocks, thus over two sessions 864 trials (12 blocks * 72 trials) were recorded.

In the *Stim* trials, participants received randomly either a TMS or a TES pulse at a randomized latency as they performed reaching movements. TMS and TES were delivered at 4 latencies in the pro and anti-blocks, 3 of which were common for both reach types across all participants: 60, 80 and 100 ms after target presentation (**Figure 10D**). The “voluntary” latencies for pro- and anti-reaches were calculated as in experiment 1; based on the median onset times of pectoralis major muscle EMG in session one. We reduced the number of stimulation latencies and excluded the *noStim* condition in experiment 2 because adding TES to the experiment design significantly increased the number of trials and lengthened the duration of the experiment. The *noStim* trials were identical to experiment 1; participants were not given brain stimulation while doing the reaching movement. To summarize, the *Stim* trials had 32 conditions: 4 latencies (60, 80, 100, voluntary) x 2 (left and right target) x 2 (pro and anti-reach) x 2 (TES and TMS). The *noStim* trials had 4 conditions: 2 (left and right target) x 2 (pro and anti-reach). The *Stim* and *noStim* conditions had 24 trials each. Across the 12 stimulation blocks there were 768 *Stim* trials, 96 *noStim* trials. Considering the transient discomfort that accompanies TES, a constraint was imposed in the trial sequence throughout the experiment to prevent more than two consecutive TES trials.

### Data Analysis

#### Motor Evoked Potentials

The primary variable of interest in this study was the amplitude of the motor evoked potentials (MEP) produced by the brain stimulation. We used MEP amplitudes to quantify the influence of the visual target on the responsiveness of the corticospinal tract. Specifically, we analyzed how the amplitude of MEPs in the four muscles varied as a function of stimulation latency and visual target direction. To calculate the MEP in each stimulation trial, a 150 ms window (-50 ms to 100 ms with respect to the stimulation onset) was extracted from all four muscle EMGs. The amplitude of a MEP was then calculated on a trial-by-trial basis by taking the mean of the rectified EMG signal within a specific MEP window. The trial-by-trial MEP amplitudes were then averaged to get the mean value per condition. The MEP window for each muscle varied slightly across participants. For every participant, the start and end latency of the MEP window in each muscle was assigned by visual inspection of the average MEP trace (See green dotted box in **Figure 3A** for example MEP window). To compare across participants, we normalized the MEP amplitudes to a standard. For the first experiment, MEP amplitudes in each “*Stim* trial” condition were expressed as a percentage of the average MEP size in the *noTarget* trials. For the second experiment, average MEP amplitudes in each “*Stim* trial” condition were expressed as a percentage of the average MEP size in the no movement trials in session one. It is worth noting that the main results in this study were identical when we used other methods of MEP quantification such as the peak-to-peak amplitude of the averaged waveform.

For experiment two involving TMS and TES, the comparison of MEP onset latency was critical to indicate a different site of corticospinal activation. We used a double-blind method to determine the onset latency of the Motor Evoked Potentials from TMS and TES. We used the trials from the first block (40 trials total: 20 TMS, 20 TES) in the experiment where participants were not reaching and just holding a steady background contraction. To determine the onset latency, a custom MATLAB script generated plots with the mean MEPs overlaid on the individual MEP traces for each participant. No markers that could identify the stimulation type or the participant were present in the figures. Forty such figures were generated: 20 subjects by 2 stimulation types (TMS and TES). These figures were then randomised and presented to a second researcher who did not take part in the above steps. The custom MATLAB script allowed interactive clicking on waveform plots and digitising of data points selected. The researcher then carefully observed the MEP waveform and used mouse clicks to identify the point at which the wave first became distinguishable from the baseline. This point was identified as the onset latency of the MEP. This process was then repeated for each of the 40 figures.

#### Kinematic Analysis

Acceleration data were converted to velocity using the cumulative integral function (cumtrapz) in MATLAB. For a given reach, reaction time (RT) was determined by first identifying the point of peak velocity and then going backward in time to the nearest point when velocity reached 5 % of peak. Only the first peak which corresponded to the initial arm movement was considered for picking the peak velocity. We excluded trials in which reaction times fell outside the range of 140-500 ms. Reach direction was identified by checking the relative sign of the velocity curve (positive indicates a left reach and negative indicates right reach) at peak velocity.

#### Express visuomotor Response and EMG onset Detection

We detected the presence of express visuomotor responses in participants using the Receiver Operator Characteristic (ROC) analysis described in Contemori et al. 2020. This method enabled us to determine a discrimination time when the laterality of the target could be reliably differentiated from EMG alone by an ideal observer. Briefly, for each reach condition (pro and anti) and each target location (left and right), we divided the trials based on reaction time into a fast trial group and a slow trial group. We then grouped the fastest trials for the left and right target together and similarly grouped the slowest trials. We then ran the ROC analysis on every sample (1 ms) between 100 ms before to 300 ms after target presentation. For each time point the ROC analysis yields the Area Under the ROC Curve (AUC), which has values ranging from 0 to 1. A value of 0.5 indicates that an ideal observer will have random chance at distinguishing the target location from the EMG, whereas a value of 1 indicates perfect discriminability and 0 indicates perfect non-discriminability. The ROC analysis output a curve with AUC on the y-axis and time along the x-axis. We defined discrimination time as the time after target presentation when the value of the ROC curve exceeded 0.67 and remained over this value for at least 15 out of the next 20 ms.

A participant was considered as potential express response positive if the discrimination time for both fast and slow trials were within the express response time window of 70-120 ms. If the discrimination time for both fast and slow trials were within the express response time window, then we checked whether the discrimination time was time-locked to the target-onset or covaried with the manual reaction time. To determine this, we fit a line to the two data points generated by plotting the median reaction time (y-axis) and discrimination time (x-axis) for the slowest and fastest trials. Perfect covariance between discrimination time and manual reaction time would mean that EMG activity is time locked to onset of hand motion and not the target appearance; this would be represented by a slope of 45°. If the slope is 90°, then discrimination time is fully independent of the manual reaction time (slowest and fastest trials have same discrimination time), and EMG is time-locked to the peripheral target appearance. We classified a participant as express response positive if the absolute slope exceeded 67.5°. For a positive participant, we re-ran the ROC analysis for all the trials (fast and slow combined) to determine the discrimination time for each tested condition.

We also performed single-trial analysis to detect the onset of EMG activity on each individual reach using the *detrended-integrated* signal method described in Contemori et al. 2022. Briefly, for each correct trial, we took a time window -100 ms before to 300 ms after the onset of the target for analysis. We then subtracted the average muscle activity in the baseline period (-100 to 70 ms w.r.t target presentation) from the entire EMG signal and then computed the integral of this signal. We then detrended this signal by subtracting the linear regression of the background period from the signal to obtain a detrended-integrated signal. We defined a candidate EMG response time as the time point when the detrended-integrated signal diverged five standard deviations from the average baseline period activity of the detrended-integrated signal. If the signal was five standard deviations above the baseline activity, then it was classified as EMG activation and if the signal was five standard deviations below the baseline activity, then it was classified as EMG inhibition. To find the initial deflection point in the detrended-integrated signal we performed a linear regression analysis around the candidate response time. For muscle activation, we fitted a linear trendline to the data between the last valley before and the first peak after the candidate response time. For muscle inhibition, we fitted a linear trendline to the data between the last peak before and the first valley after the candidate response time. Finally, the EMG activity onset time was defined as the time point at which the linear trendline met the line representing the average baseline period activity of the detrended-integrated signal.

#### Statistical analysis

We employed frequentist statistical analyses using JASP (JASP Team 2022, Version 0.16.2) and GraphPad Prism (Version 9.4.0, GraphPad Software, San Diego, California US). Our analysis models focused on how the amplitudes of the motor evoked potentials (MEPs) were impacted by the interactions between the latency of stimulation and the location of the visual target. For the first experiment, we ran repeated-measures ANOVAs on MEPs, with the target location and stimulation latencies as the repeated measures. For the second experiment, we ran repeated-measures ANOVAs on the MEP difference between left and right targets, with stimulation type (TMS v TES) and stimulation latencies (60, 70, 80, 90, 100, and *voluntary*) as the repeated measures factors. For every muscle, ANOVA was run separately for pro- and anti-reach trials. Greenhouse-Geisser correction was applied whenever Mauchly’s test of sphericity was violated. When there was a significant main effect or interaction in the ANOVA results, we used Holm-Šídák test for post-hoc comparison. For all tests, statistical significance was set at P < 0.05. We inspected the QQ plots to check whether data followed a normal distribution. All distributions were approximately normal and parametric statistics were used throughout.

